# A microRNA atlas of the human prefrontal cortex across the adult lifespan

**DOI:** 10.64898/2026.05.27.728239

**Authors:** Irais Erendira Valenzuela-Arzeta, Pourya Naderi Yeganeh, Anupama Rai, Jane Banahan, Artemis Iatrou, Leinal Sejour, Madison Uyemura, Isaac H. Solomon, Nikolaos P Daskalakis, Ioannis Vlachos, Winston Hide, Frank J. Slack, Maria Mavrikaki

## Abstract

Aging of the human brain is characterized by widespread changes in gene expression regulated in part by microRNAs (miRNAs). We present a lifespan miRNA atlas of the human dorsolateral prefrontal cortex generated from small RNA sequencing of 113 postmortem samples spanning 18 to 100 years of age. Differential expression analysis revealed progressive age-associated remodeling of miRNA expression, with the strongest differences observed between old and young individuals. Among the significantly altered miRNAs, miR-34a-5p emerged as one of the most robustly upregulated miRNAs in the aged cortex, alongside additional aging-associated miRNAs including miR-155-5p, miR-132-3p, miR-212-3p, miR-449a, and members of the miR-302 family. This atlas provides a resource for investigating miRNA dysregulation and small RNA regulatory networks during human cortical aging.

## Introduction

Aging is associated with molecular and functional changes in the human brain that contribute to cognitive decline and increased susceptibility to neurodegenerative disease^1,2^. The dorsolateral prefrontal cortex (DLPFC), a region critical for executive function and working memory, is particularly vulnerable to aging-related remodeling and neurodegenration^3–5^. Recent transcriptomic studies including ours indicate that cortical aging is spatially structured and cell-type-specific rather than uniform across tissue^1,2,6^.

MicroRNAs (miRNAs) are small non-coding RNAs that regulate gene expression post-transcriptionally by targeting messenger RNAs^7^. In the brain, they play essential roles in neuronal differentiation, synaptic function, and stress responses^8–10^, and their dysregulation has been linked to neuroinflammation and neurodegeneration^11^. Previous studies have reported age-associated miRNA changes in human brain^2,6,9,11–15^, with the miR-34 family emerging as a conserved regulator of aging and neuronal stress pathways^9,11,14,15^. However, the landscape of lifespan-associated miRNA expression changes in the human cortex remains incompletely characterized.

Here, we profiled miRNA expression across the adult lifespan in the human DLPFC using small RNA sequencing from 113 postmortem samples spanning 18 to 100 years of age. We identified progressive age-associated remodeling of miRNA expression, with the strongest transcriptomic differences observed between old and young individuals. Among the significantly altered miRNAs, miR-34a-5p emerged as one of the most robustly upregulated miRNAs in the aged cortex. Together, our findings provide a lifespan miRNA atlas of the human DLPFC and establish a resource for investigating small RNA regulatory networks during human brain aging.

## Results

To characterize age-associated miRNA expression changes in the human DLPFC, we performed small RNA sequencing on 113 postmortem samples spanning the adult lifespan (18-100 years; Fig. 1A) and categorized them as young (<40 years), middle-aged (41–69 years), and old (≥70 years). Differential expression analysis revealed that age-dependent miRNA expression changes were most pronounced in the old versus young comparison (Fig. 1B). In contrast, the middle aged versus young and old versus middle aged comparisons exhibited fewer significant alterations, supporting a model of progressive transcriptomic remodeling during aging rather than abrupt shifts, consistent with emerging transcriptomic atlases of aging tissues^1,16,17^.

**Figure 1.**
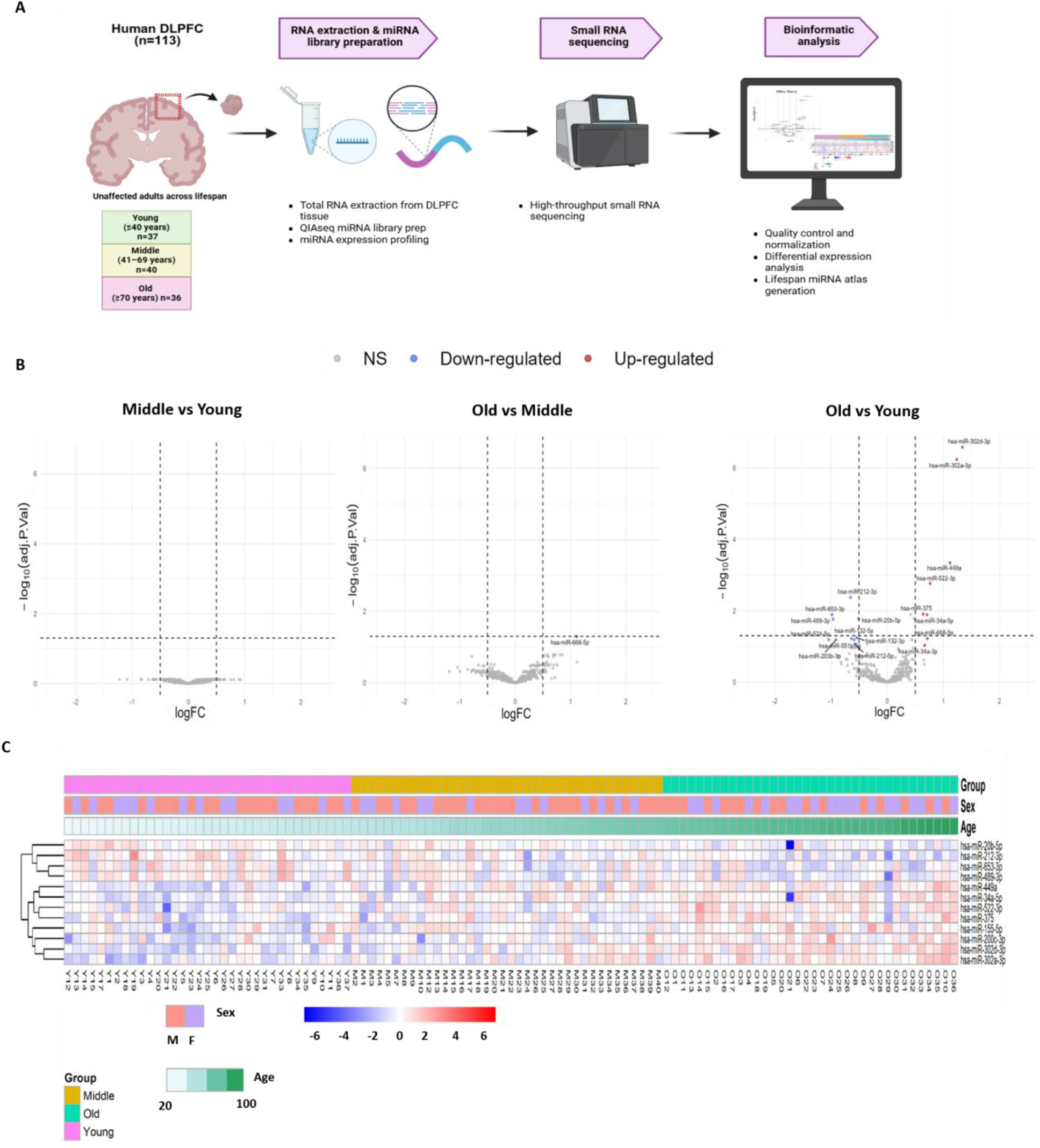
Study design and age-associated miRNA expression changes in the human dorsolateral prefrontal cortex (DLPFC). (A) Overview of the experimental design. Small RNA sequencing was performed on postmortem human DLPFC samples across the adult lifespan (18-100-years old). (B) Volcano plots showing differential miRNA expression for middle aged versus young, old versus middle aged, and old versus young comparisons. miR-34a-5p is highlighted among significantly altered miRNAs. (C) Heatmap of differentially expressed miRNAs across all samples. Samples are placed on an ascending age order (18-100-years old). Expression values are shown as row-scaled z-scores with hierarchical clustering of miRNAs.

Several miRNAs exhibited marked differential expression in old versus young comparisons, including miR-34a-5p, miR-302d-3p, miR-302a-3p, miR-155-5p, miR-449a, miR-522-3p, miR-132-3p and miR-212-3p, among others^8,18^ (Fig. 1B). The presence of multiple brain-related and developmentally implicated miRNAs among the top differentially expressed candidates supports the biological robustness of the dataset and establishes it as a resource for studying age-associated miRNA regulatory networks in the human cortex^19^.

Unsupervised hierarchical clustering of differentially expressed miRNAs (FDR < 0.05) further demonstrated coordinated expression changes across age groups (Fig. 1C). Samples ordered by chronological age revealed a gradual shift in miRNA expression patterns, with older individuals exhibiting coherent up- and down-regulation signatures. Among the significantly altered miRNAs, miR-34a-5p (miR-34a) emerged as one of the most robustly upregulated miRNAs in aged individuals (logFC = 0.712, FDR = 0.0129) consistent with experimental models of prefrontal aging demonstrating miR-34a overexpression and associated neuronal apoptosis^14,16,20^. Inspection of individual samples demonstrated a consistent elevation of miR-34a expression across the majority of old subjects (Fig. 1C). Clustering analysis also demonstrated that miR-34a and miR-449a, members of the same family with the same seed sequence, exhibit highly correlated expression trajectories across the lifespan. This reproducible age-associated increase positions miR-34a as a strong candidate for downstream analyses^14,20^.

## Discussion

Our findings identify miR-34a as a reproducibly upregulated miRNA in the aging human DLPFC and expand previous observations linking the miR-34 family to aging and neuronal stress responses^15,21^. While prior studies have implicated miR-34a in aging-associated pathways, including cellular stress responses, apoptosis, and neurodegeneration, lifespan-wide characterization of miRNA expression changes in the human cortex has remained limited. Here, using small RNA sequencing across 113 human DLPFC samples spanning adulthood, we provide a resource defining age-associated miRNA remodeling in the human cortex.

Our analysis revealed that age-associated miRNA changes were most pronounced between old and young individuals, whereas middle-aged comparisons demonstrated fewer alterations, supporting a model of progressive transcriptomic remodeling during aging rather than abrupt molecular transitions. Among aging-associated miRNAs, miR-34a was one of the most robustly upregulated candidates. Increased miR-34a expression has previously been linked to neuronal stress responses, apoptosis, and aging-related molecular pathways, supporting its potential role as a conserved regulator of brain aging^14,15,21^

Beyond miR-34a, we identified additional aging-associated miRNAs, including miR-155-5p, miR-132-3p, miR-212-3p, and members of the miR-302 family which have also been previously implicated in aging processes^12,13,22^. Interestingly, only miR-668-5p, which has also been linked to senescence^23^, was significantly upregulated in the brains of old individuals compared to the brains of middle-aged individuals as well as the brains of young individuals. Notably, several of these miRNAs are key regulators in Alzheimer’s disease, suggesting that aging-associated miRNA changes may represent early molecular events contributing to disease vulnerability^2,12,13,24^

Clustering analysis demonstrated that miR-34a and miR-449a exhibit highly correlated expression trajectories across the lifespan^22^. Both miRNAs were significantly upregulated in the brains of old individuals compared to middle-aged and young groups. These findings align with previous studies showing that miR-34a acts synergistically to miR-449a on autophagy^23^, suggesting a coordinated synergistic regulation of autophagy-related pathways in the aging brain.

Taken together, this study provides a lifespan miRNA atlas of the human DLPFC and establishes a resource for investigating how small RNA regulatory networks evolve during human brain aging. Defining age-associated miRNA expression changes across the lifespan may contribute to understanding molecular mechanisms underlying normal brain aging and vulnerability to neurodegenerative disease.

## Acknowledgements

This study was supported by NIH grants to MM (5R01AG079799) and FS (R01AG058816 and R01AG082093). We thank the NIH NeuroBioBank (University of Maryland, University of Miami, and Harvard Brain Tissue Resource Center; https://neurobiobank.nih.gov/) and the Human Brain Collection Core of the NIMH Intramural Research Program (http://www.nimh.nih.gov/hbcc) for providing the tissues analyzed in this study. We also thank Stefano Marenco, Sabina Berretta, Alexandra LeFerve, and David Davis for assistance with the identification of appropriate tissues, and the Harvard Biopolymers Core for performing quality control on our samples. The authors acknowledge the use of the BIDMC Ithaca High Performance Computing Cluster of the Spatial Technologies Unit (RRID:SCR_024905), which supported a portion of the analyses.

## Material and Methods

### Human Postmortem Brain Tissues

We analyzed frozen frontal cortex tissue blocks (Brodmann area 8, dorsolateral prefrontal cortex, DLPFC) from a total of 113 donors obtained through the NIH NeuroBioBank (University of Maryland, University of Miami, and Harvard Brain Tissue Resource Center, HBTRC) and the NIMH Human Brain Collection Core (HBCC). The cohort included 66 males and 47 females, across the adult lifespan ranging from 18 to 100 years (see Table S1). Based on age, donors were categorized into three groups: young (18-40 years, n = 37), middle-aged (41–69 years, n = 40), and older adults (70-100 years, n = 36) according to our previous studies^6,31^. Selection criteria for specimens from the NIH NeuroBioBank required the absence of a reported clinical brain diagnosis (unaffected controls), a postmortem interval (PMI) shorter than 48 hours, and an RNA Integrity Number (RIN) above 7. Cases with neurological or psychiatric disorders, as well as those with HIV, or hepatitis C virus (HEPC), were excluded for biosafety reasons. Cases with COVID-19 were excluded from this cohort as we previously shown that severe COVID-19 is associated with transcriptomic changes of aging^25^.

### RNA Extraction and Quality Assessment

Total RNA was extracted from brain tissue using the TRIzol reagent (Thermo Fisher Scientific) and phase separation. RNA quality was evaluated at the Biopolymers Facility (BPF Genomics and Core Facility, Harvard Medical School) using the Agilent TapeStation 4200 Instrument.Samples with RNA Integrity Number (RIN) below 4.0 (as measured in our lab) were not included in the study.

### Small RNA library construction

Small RNA libraries were prepared from total RNA using the QIAseq® miRNA UDI Library Kit (QIAGEN, Hilden, Germany), following the manufacturer’s protocol. Briefly, adapters were sequentially ligated to the 3′ and 5′ ends of mature miRNAs, followed by reverse transcription with primers containing unique molecular identifiers (UMIs). cDNA products were purified with bead-based cleanup and subsequently amplified by 16 cycles of PCR using unique dual indexes (UDIs) to enable multiplexed sequencing. Final libraries were assessed for size distribution (∼200 bp) and concentration using an Agilent TapeStation 4200 Instrument and Qubit fluorometer (ThermoFisher Scientific). Libraries were pooled into two pools and sequenced on an Illumina NovaSeq platform using paired-end 150 bp reads (Illumina, San Diego, CA, USA).

### Small RNA-Seq data processing

Raw small RNA count matrices for the combined imported and harmonized across two batches of sequencing. Lowly expressed RNAs were identified and excluded based counts per million (CPM) reads, (at least 1 CPM across a minimum of 20% of the samples). Age strata were defined in metadata as Young (<40 years), Middle (40–69 years), and Old (≥70 years), with ordered factor levels Young/Middle/Old. Data cleaning and normalization were performed using utilities provided in the edgeR and limma packages.

Normalized counts for PCA were obtained via DESeq2 size-factor estimation, followed by log-transformation and PCA on samples. PCA was visualized against biological/technical covariates (Sex, Age, RIN, Library prep) to identify confounding effects and outliers. A single library-prep batch was identified as an outlier and removed prior to final analyses; corresponding samples were excluded from both metadata and count matrices. To quantify the variance explained by covariates for downstream inclusion in linear models, we used variancePartition using a mixed-effects model random effects for categorical variables Group, Library prep, and Sex, and fixed effect for the continuous RIN (as suggested in the package manual)^26^.

### Differential expression analysis

Differential expression analysis was performed between different age groups accounting for confounding covariates including RIN, age, sex, and library preparation. Filtered counts were modeled using limma-voom with a design matrix including Group (age strata), RIN, library preparation, sex, and Batch. We contrasted Middle vs Young, Old vs Young, and Old vs Middle. P-values for differential expression were estimated using empirical Bayes moderation (eBayes). Differential expression p-values were adjusted for multiple testing using Benjamini-Hochberg FDR. Significant differential expression events were determined using an FDR threshold of 0.05. To generate miRNA-specific outputs, we filtered the DE results to annotated miRNAs and recalculated the FDR values.

